# Assessing seabird and shorebird vulnerability to offshore wind farms in Brazil: A decision-support framework for environmental licensing

**DOI:** 10.64898/2026.09.29.755444

**Authors:** Natascha Horn, Guilherme Tavares Nunes

## Abstract

Environmental concerns and the need to mitigate the effects of climate change have driven the adoption of renewable energy sources, such as offshore wind farms (OWFs), making it essential to assess their overlap with seabird habitats to estimate potential impacts. This study is the first to estimate the vulnerability of coastal and seabirds to OWFs at a national scale by integrating biological trait data from the literature, spatial occurrence records from citizen science platforms, and expert knowledge (elicited from 23 specialists) into established quantitative risk frameworks. Collision risk and susceptibility scores were calculated at the family and subfamily levels, as well as disturbance and displacement indices by species and susceptibility by state. The groups with the highest collision risk and susceptibility were Fregatidae (0.354; 0.486), Sulidae (0.199; 0.419), Sterninae (0.190; 0.384) and Diomedeidae (0.190; 0.416). The extreme risk for Fregatidae is driven by their specific flight characteristics, with impacts concentrated in the highly frequented southeast and northeast regions. Regarding disturbance and displacement, *Limnodromus griseus* exhibited the highest national score (MMA: 0.098), while *Diomedea dabbenena* had the highest global score (IUCN: 0.092), highlighting the critical need to integrate both regional and global conservation statuses. Spatially, the state exhibiting the highest overall vulnerability to OWF impacts was Rio Grande do Sul (23.78; 10.69), a critical hotspot characterized by high species richness and oceanographic productivity. These findings establish an indispensable national baseline for Marine Spatial Planning, reinforcing the need for targeted pre-installation assessments, the identification of sentinel species, and spatially differentiated management during environmental licensing processes.

**Graphical Abstract:** 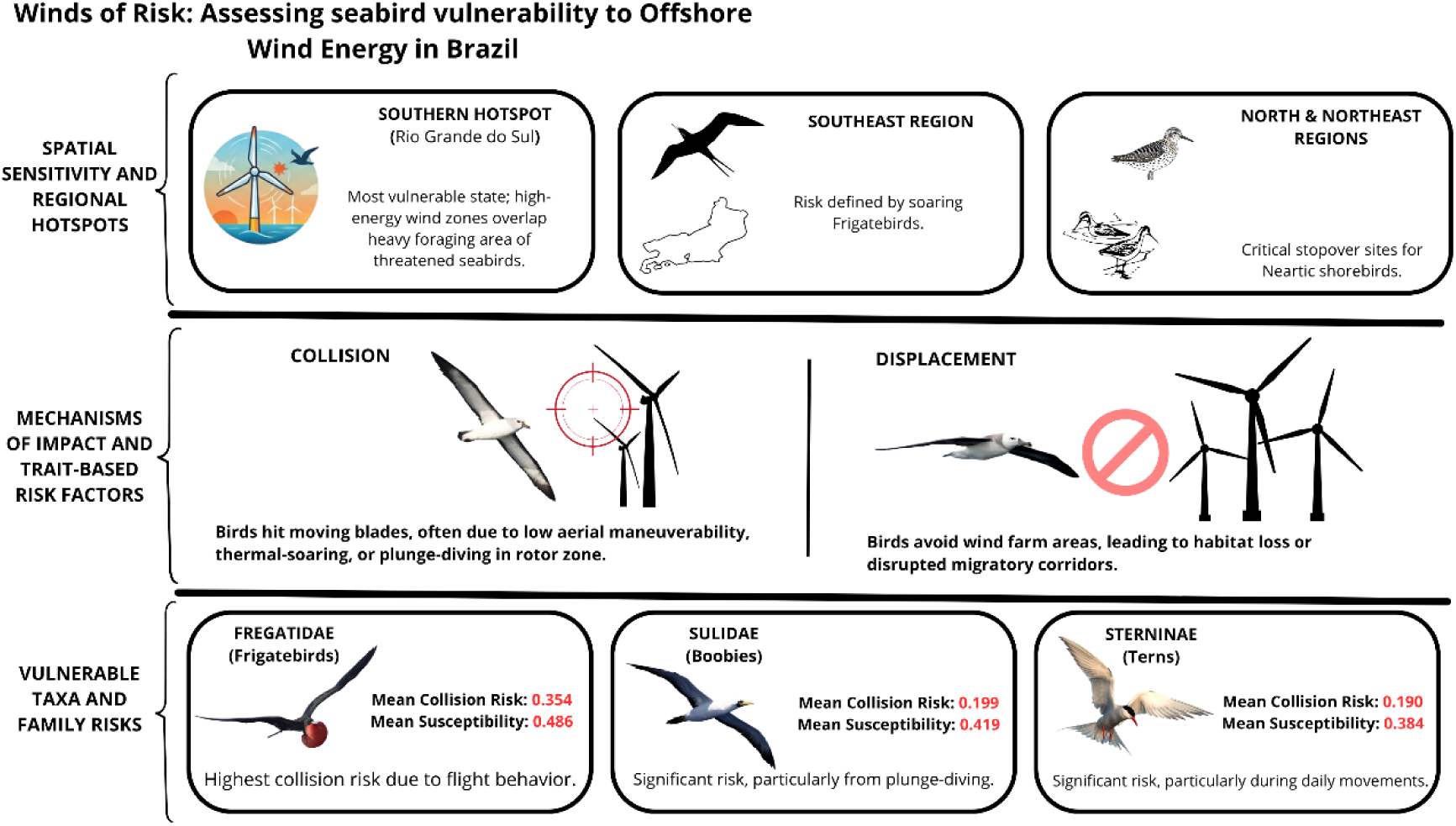

## 1. Introduction

The global expansion of offshore wind farms (OWFs) has introduced a new frontier of environmental challenges for marine biodiversity (Watson et al., 2024a). Among the most vulnerable taxa are seabirds, whose life-history traits such as long-distance migrations, low reproductive rates, and reliance on specific marine habitats make them particularly susceptible to anthropogenic disturbances (Bradbury et al., 2014; Schreiber, 2002; VanCompernolle et al., 2026). Although OWFs are widely recognized as a promising pathway toward decarbonization and energy security, their rapid deployment in regions lacking ecological data raises concerns about unintended ecological impacts (Laurino et al., 2025; Morant et al., 2025). The lack of biodiversity baselines and species-specific sensitivity assessments hinders the development of effective mitigation strategies and environmentally sound marine spatial planning (Abramic et al., 2022; White et al., 2024). Seabirds serve as sentinels of ocean health and indicators of ecosystem change, making their inclusion in impact assessments essential for aligning offshore wind development with ecosystem-based management (Garthe et al., 2023). This task becomes particularly complex and urgent in countries with extensive coastlines and high seabird diversity, such as Brazil (Bugoni et al., 2022).

Brazil hosts one of the longest coastlines in the world, stretching over 8,000 km and spanning approximately 38 degrees of latitude. This extensive and heterogeneous marine landscape supports a rich assemblage of seabirds, comprising nearly one-third (∼100 species) of the approximately 350 seabird species recognized globally (Bugoni et al., 2022). Although a portion of these are vagrant species with occasional records of occurrence (Pacheco et al., 2021), Brazil’s coastal and oceanic zones are critical for both tropical and temperate seabirds, serving as breeding, foraging, and migratory habitats. The country’s Exclusive Economic Zone (EEZ) functions as a wintering area for both austral and boreal migrants, as well as a breeding ground for resident species that nest on neritic and oceanic islands (Mancini et al., 2016).

In recent years, Brazil has experienced a rapid surge in offshore wind energy proposals (Bugoni et al., 2022). According to the Brazilian Institute of the Environment and Renewable Natural Resources (IBAMA), 16,178 turbines distributed across 104 projects are currently under environmental licensing, with a combined projected capacity of 247,354 MW (IBAMA, 2025a). Notably, most of these proposed projects overlap with key foraging and migratory areas for seabirds, highlighting the urgent need for robust baseline data to inform environmental impact assessments and ensure that licensing decisions are ecologically sound (Bugoni et al., 2022).

To understand the magnitude of these potential impacts, it is essential to consider the scale and technical characteristics of modern offshore wind farms. Unlike land-based structures, offshore projects utilize massive wind turbines with nominal capacities frequently ranging from 10 to 15 MW, rotor diameters exceeding 200 meters, and total heights that can surpass 250 meters. Depending on the depth of the continental shelf, these turbines are deployed using fixed foundations (such as monopiles or jackets, in waters up to 50 to 60 meters deep) or mounted on floating structures in deeper areas. Arranged in spatial layouts spanning tens to hundreds square kilometers, these parks introduce unprecedented vertical and dynamic elements into the marine environment. It is precisely these physical and operational dimensions that begin to interact with the local ecosystem (EPE, 2020; Hernandez et al., 2021).

Offshore wind turbines can affect seabirds through a range of mechanisms that are often species- and context-specific (Bailey et al., 2014). The two primary types of impact are collision, when birds are unable to avoid moving blades and suffer injury or mortality, and displacement, which occurs when individuals alter their spatial behavior, either by avoiding areas entirely or modifying their movement and foraging patterns in response to turbine presence (King, 2019; Perrow, 2019). These responses may lead to the loss of foraging grounds, disruption of movement corridors, increased energetic demands, or even ecosystem-level trophic shifts. Displacement effects can manifest at different spatial scales, from broad avoidance of entire wind farms (macro scale) to alterations in flight paths within the complex (meso scale), or fine-scale behavioral changes near turbines (micro scale) (Thaxter et al., 2017). In some situations, seabirds may also be attracted to turbine structures, using them as roosting sites or exploiting changes in local prey availability (Dierschke et al., 2016; Vanermen et al., 2015).

Despite growing awareness of these potential impacts, most of the available knowledge stems from studies conducted in the Northern Hemisphere, where monitoring infrastructure and regulatory frameworks are more developed (Agudelo et al., 2021; Watson et al., 2024b). In contrast, empirical data from tropical and subtropical systems such as those found along the Brazilian coast remain scarce, limiting the ability to detect, attribute, or forecast the consequences of offshore wind development under different ecological conditions (Bugoni et al., 2022).

The logistical challenge of detecting seabird collisions at sea, combined with the need for detailed pre-installation data to assess behavioral displacement, represents a major barrier to effective impact assessment. Moreover, responses to OWFs can differ significantly even among closely related species, depending on their ecological traits, prey distribution, and the spatial configuration of turbines (Croll et al., 2022; Pollock et al., 2024). These challenges are further compounded by the cumulative effects of associated infrastructure such as substations, undersea cables, and coastal transmission lines, which must be considered alongside turbine-related impacts (Farr et al., 2021; Goodale and Milman, 2019). Together, these knowledge gaps underscore the urgency of generating robust baseline data on seabird spatial ecology and sensitivity in Brazilian waters to support evidence-based licensing and long-term monitoring (Masden et al., 2010).

Several quantitative approaches have been proposed to assess seabird vulnerability to OWFs (Cheng et al., 2026; Fauchald et al., 2024). These include collision and displacement risk indices based on behavioral, ecological, and conservation attributes (Croll et al., 2022; Furness et al., 2016, 2013; Garthe and Hüppop, 2004; Reid et al., 2023), as well as ecological niche modeling to predict spatial overlap with planned infrastructure (Lemos et al., 2023; Morant et al., 2025). Ideally, a comprehensive vulnerability assessment would rely on an empirical gold-standard dataset. This would encompass nationwide, long-term tracking data across dozens of sentinel species to capture fine-scale dynamic interactions with the airspace, paired with automated land-based monitoring and dense, multi-year at-sea surveys capable of resolving both spatial distribution and macro-seasonal variations along the entire coastline. However, acquiring such standardized, large-scale empirical data requires multi-decadal investments and faces profound logistical constraints in the Global South. The implementation of these robust predictive frameworks demands extensive, high-resolution empirical data on pelagic distribution, precise flight heights, and *in situ* avoidance behaviors (Kelsey et al., 2025). This data deficiency is highly concerning given the rapid expansion of offshore wind development interests in these areas, which creates a critical gap in the scientific subsidies required for informed decision-making during the environmental licensing process regarding seabirds. In the absence of this ideal dataset, alternative analytical frameworks that synthesize multi-source data become paramount to provide immediate, scientifically grounded support for current environmental licensing demands.

To overcome these limitations, expert-based approaches provide an effective alternative for generating structured, repeatable estimates of species sensitivity in data-poor regions (Hemming et al., 2018; Martin et al., 2012). In Brazil, although refined behavioral datasets are limited, researchers have accumulated extensive field knowledge through systematic seabird censuses conducted during fisheries monitoring programs, targeted research initiatives, and logistical cruises to coastal and oceanic islands (Franz et al., 2021; Lemos et al., 2023; Valls et al., 2023). These opportunistic yet methodologically consistent efforts offer valuable insights into key biological attributes such as species distribution, flight behavior, and spatial use of marine habitats. Critically, this experiential knowledge acquired through repeated *in situ* observation can serve as a robust foundation for estimating the parameters used in vulnerability indices. In contexts where formal literature is scarce but development pressure from OWFs is mounting, integrating this type of field-derived expertise becomes essential for producing informed, ecologically meaningful assessments (Drescher et al., 2013; Lemos et al., 2023).

This study represents a nationwide assessment that integrates multiple layers of data to fill knowledge gaps regarding the impact of offshore wind energy in Brazil. The approach combined biological data from the literature with empirical knowledge from 23 experienced field ornithologists, obtained through structured questionnaires, to generate a synthesis of seabird ecology. Additionally, the use of regional species lists, and citizen science data allowed for the spatialization of vulnerability patterns along the 8,000 km of the Brazilian coastline. This multi-source approach supports decision-making in environmental licensing and adaptive management in the sector. Furthermore, the scope of the study is aligned with national conservation priorities, such as the IBAMA Official Guidelines (2020) and the National Action Plans (PLANACAP and PAN Aves Marinhas) and converges with international agreements such as the Agreement on the Conservation of Albatrosses and Petrels (ACAP) and the Convention on Migratory Species (CMS).

## 2. Methodology

### 2.1 Data collection

The study area encompasses the entire Brazilian coastline, extending from the state of Amapá in the north (4.2°N) to Rio Grande do Sul in the south (33.4°S), and includes 17 coastal states and 279 municipalities facing the sea (Table S1; Figure 1). Brazil’s coastline forms a continuous stretch of over 8,000 km in length, characterized by a wide variety of coastal landscapes such as dunes, islands, reefs, rocky shores, bays, estuaries, mangroves and sea cliffs (MMA Ordinance No. 34/2021; IBGE).

**Figure 1.**
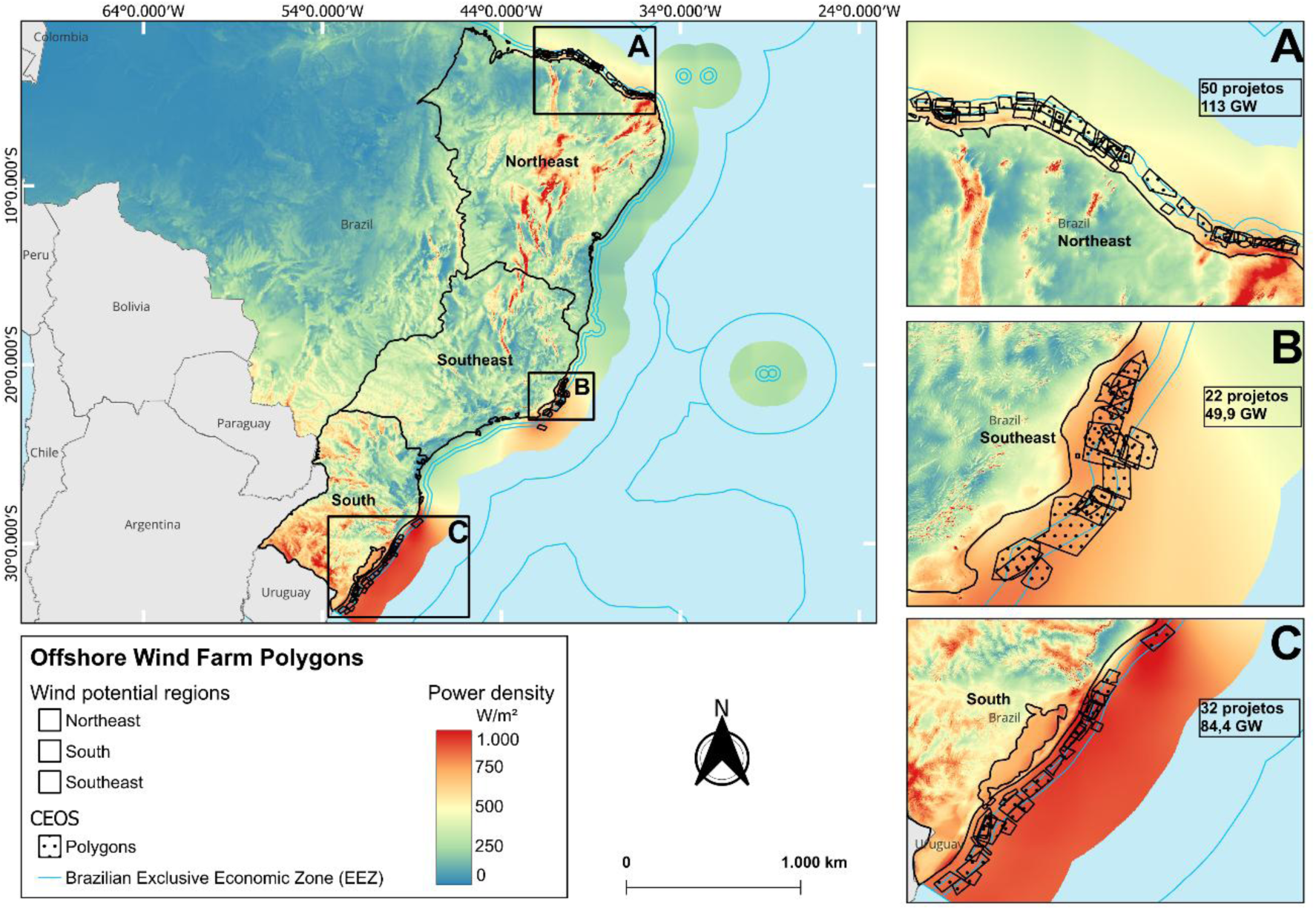
Study area along the Brazilian coast, highlighting the three macro-areas with highest wind energy potential (A – Northeast region, B – Southeast region, C – South region) and the location of polygons in the licensing process by IBAMA.

A biological database was compiled using a questionnaire (Appendix S1) completed by specialists for each seabird group covering all species with regular occurrence in Brazilian territory, following (Pacheco et al., 2021) and (Nunes et al., 2023). Species-level data on biological traits and habitat use were collected, following the same parameters described by (Garthe and Hüppop, 2004), and later adopted by (Furness et al., 2013) and (Reid et al., 2023) (Table 1). Based on this dataset, species-specific scores were calculated for collision risk, disturbance/displacement risk (as described by (Furness et al., 2013), and overall susceptibility (as described by (Reid et al., 2023). In addition, species lists were generated for each coastal state in Brazil, allowing for the calculation of seabird sensitivity indices to OWFs at both the state and national levels (Franz et al., 2018; Just et al., 2018; Lima et al., 2022; Sagot-Martin et al., 2020; Silveira and Uezu, 2011).

**Table 1.** Summary of attributes used to assess seabird sensitivity to offshore wind farms in Brazil. Detailed scoring criteria are provided in the supplementary material.

| Attribute | Source | Scoring scale |
| --- | --- | --- |
| Conservation status | Brazilian Red List (MMA 2022); IUCN (2024) | 1 (Least Concern) to 5 (Critically Endangered) |
| Flight altitude | Expert survey | 1 (0%) to 6 (81–100% of time spent flying at rotor-swept altitude) |
| Flight maneuverability | Expert survey | 1 (very high) to 5 (very low) |
| Nocturnal flight activity | Expert survey | 1 (none) to 5 (intense activity) |
| Habitat specialization | Expert survey | 1 (not present in wind farm depth range) to 3 (restricted to 5–50 m depth) |
| Proportion of time flying at sea | Expert survey | 1 (0–20%) to 5 (81–100%) |
| Disturbance response | Expert survey | 1 (no reaction or attraction) to 5 (highly sensitive and avoids disturbance) |
| Confidence level of expert response | Expert self-assessment | 1 (high: based on species-specific data) to 3 (low: inferred without direct data) |

Structured questions were sent to a group of experts pre-selected from a professional network. Age, gender, or other demographics did not affect selection. Initial experts could also refer to new peers, making this a snowball sampling method. Selected participants were researchers with expertise in seabird ecology and practical experience with ship-based offshore surveys. Individual invitations were sent via email, along with preliminary information regarding participant rights and ethical guidelines, which were reiterated in the questionnaire itself. The questionnaire was administered through the online platform Google Forms, following the approval (CAAE: 76580823.9.0000.5347) of the informed consent form (Appendix S2) by the Ethics Committee of the *Universidade Federal do Rio Grande do Sul* (UFRGS). All questions were answered individually to ensure independent responses and to enhance data reliability. Participation was voluntary; respondents were free to skip any question without justification and could withdraw from the survey at any time (Box S1). All data were stored in a secure project-specific database at the UFRGS.

Thirteen avian families were assessed based on their regular occurrence and potential exposure to OWFs impacts. Although many of the assessed taxa are considered seabirds sensu stricto, the analysis also included shorebirds and other waterbirds that may use nearshore or offshore environments and could be affected by wind energy infrastructure, such as collector substations and transmission lines. For simplicity, we hereafter refer to all assessed taxa collectively as “seabirds”. These included: Charadriidae, Haematopodidae, Chionidae, Scolopacidae, Stercorariidae, Laridae (except Sterninae), Phaethontidae, Diomedeidae, Oceanitidae, Hydrobatidae, Procellariidae, Fregatidae, and Sulidae, totaling 91 species (Table S2). The subfamily Sterninae, though taxonomically nested within Laridae, was treated separately due to its distinct morphological, ecological, and behavioral traits, and the potential to be impacted by OWFs, which justify an independent sensitivity assessment. Taxonomic classification and nomenclature followed (Pacheco et al., 2021). Species classified as occasional or vagrant were excluded from the analysis due to their low ecological relevance in the context of offshore wind energy development.

The methodology adopted in this study followed the frameworks proposed by Garthe and Hüppop (2004), Furness et al. (2013), and Reid et al. (2023), aimed at categorizing the risk of interactions between seabirds and OWFs. These studies express collision and displacement risk through a combination of susceptibility scores, enabling a species-level assessment of vulnerability to OWF presence. This standardized approach allows for the reproducibility of results across different geographic regions and study systems. Seven biological and ecological attributes were used to estimate species-specific sensitivity to OWFs. These attributes include conservation status, flight altitude, flight agility (maneuverability), nocturnal flight activity, habitat specialization, percentage of time spent flying at sea, and sensitivity to disturbance caused by turbines, vessel traffic, and helicopters. A Likert-like semi-quantitative scale was assigned to each attribute to enable the integration of expert-based and literature-derived data into a standardized and repeatable scoring framework. Each attribute was scored from 1 (lowest sensitivity) to 5 or 6 (highest sensitivity), depending on the parameter, as detailed in the supplementary material (Tables S3–S11). Subsequently, these original scores were normalized to the range of 0 to 1.

### 2.2 Data handling

After collecting answers, data was organized in a spreadsheet and assigned scores for each answer. Each response received an impact score and a confidence level ranging from 1 to 3, where 1 represented high confidence and 3 represented low confidence (Table S12). The impact score of each response was then weighed according to its confidence level. This weighting was applied by multiplying the impact score by 1 for high confidence, 0.5 for medium confidence, and 0.1 for low confidence. The calculations followed the formula below:

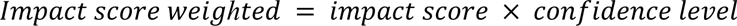

Subsequently, the mean and standard deviation were calculated for each question within each seabird family considered in the study.

### 2.3 Sensitivity indices and analytical framework

The collision risk index was calculated following the method proposed by Furness et al. (2013), considering only flight-related attributes. A higher weighting was given to flight altitude (*a*), and lower weights were assigned to flight maneuverability (*b*), proportion of time spent flying (*c*), and nocturnal flight activity (*d*):

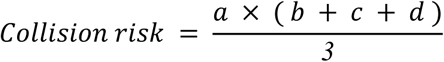

To estimate disturbance and displacement, we also followed the framework by Furness et al. (2013). The index was calculated and evaluated at the species level, considering that both national and global conservation status is assessed at this level. Therefore, we calculated a vulnerability index according to the equation below, where *f* represents the species’ sensitivity to disturbance from OWFs structures, ship traffic, and helicopters, and *g* corresponds to habitat specialization. A divisor of 10 was used to reflect the assumption that the overall impact of disturbance on bird populations is substantially lower than direct mortality from collisions.

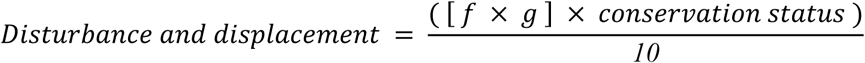

The susceptibility index was adapted from Reid et al. (2023) and was based on three attributes: flight altitude (*a*), flight maneuverability (*b*), and habitat specialization (*g*). Given the strong influence of altitude and maneuverability on seabird susceptibility to OWFs, habitat specialization was assigned a lower relative weight in the final score:

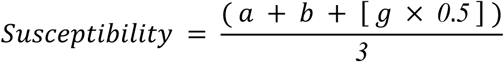

To assess seabird species richness along the Brazilian coastline, we compiled presence–absence data from the citizen science platform WikiAves (https://www.wikiaves.com.br), covering all coastal municipalities. Based on these records, we compiled species lists for each coastal state. Sensitivity indices were calculated exclusively for species confirmed to be present (Table S13).

Finally, to identify which Brazilian coastal states are most vulnerable to the impacts of OWFs, we calculated a state-level susceptibility index. For each state, we multiplied the family-level susceptibility score by the number of species from that family occurring in the state, according to WikiAves records:

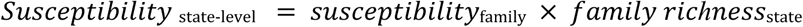

## 3. Results

### 3.1 Surveyed expert profile and experience

A total of 23 experts participated in the survey, conducted between April 10 and October 12, 2024 (Table 2; Table S14). Most respondents (70%) were affiliated with universities, followed by research institutions and non-governmental organizations (NGOs). In terms of academic qualifications, 65% hold a PhD and 35% a MSc degree. The group included university professors, graduate students, and professional researchers, all with extensive expertise in seabird ecology and marine environments. All respondents reported combining theoretical knowledge with practical field experience. Nearly all had at-sea field experience, with reported time ranging from 100 to >7,000 hours. Frequently cited regions of activity included the southern and southeastern Brazilian coast, oceanic islands, and, in some cases, international waters such as the South Atlantic and Falkland Islands.

**Table 2.**
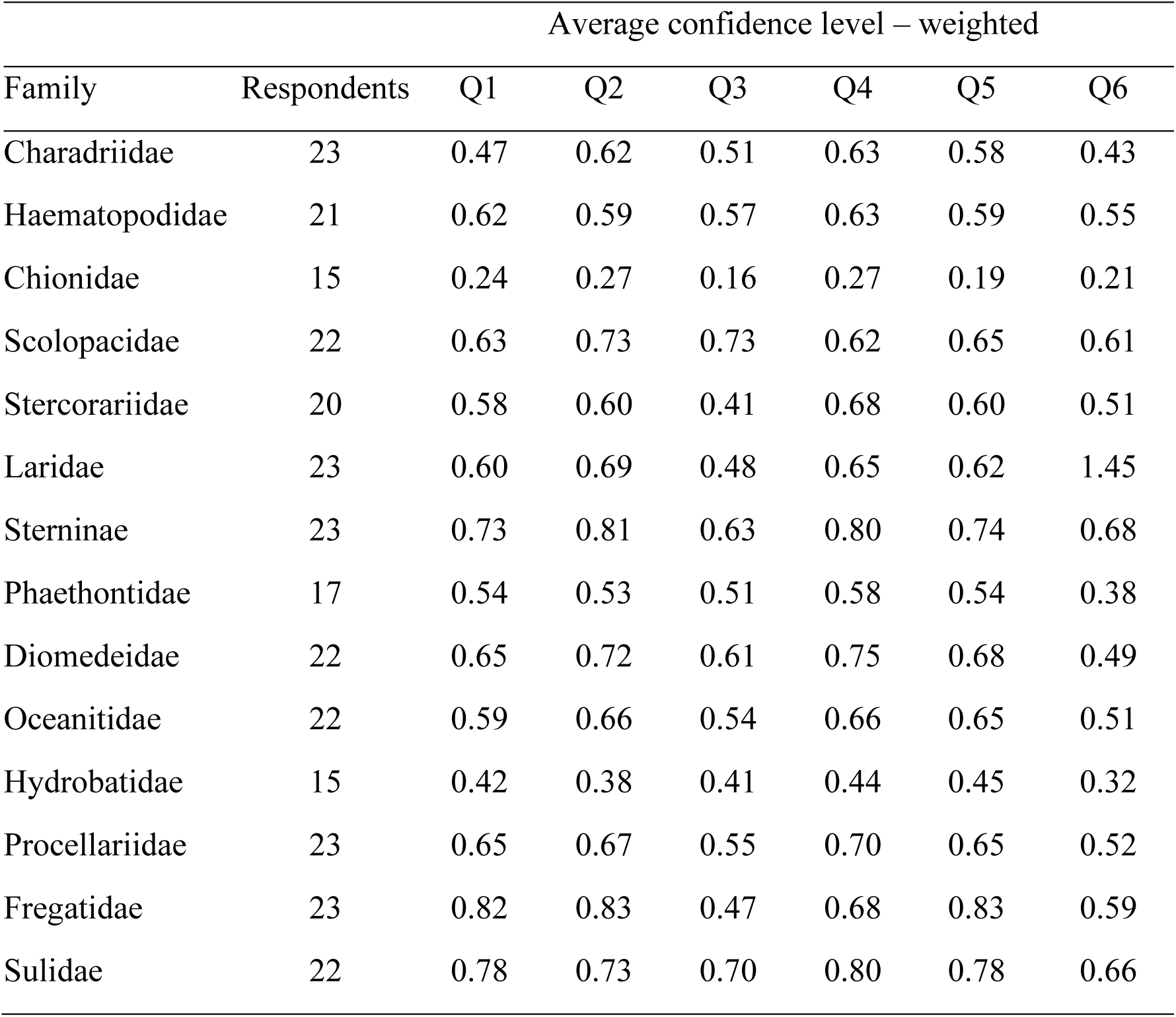
Number of respondents per (sub)family and average confidence level for each question (Q1-Q6).

### 3.2 Taxon-level vulnerability to OWFs

The families with the highest national-level mean collision risk were Fregatidae (0.354), Sulidae (0.199), and Sterninae (0.190), while the lowest were Chionidae (0.017), Hydrobatidae (0.054), and Haematopodidae (0.081) (Table 3, Figure 2). Regarding susceptibility to OWFs, the highest mean scores were recorded for Fregatidae (0.486), Sulidae (0.419), and Diomedeidae (0.416), and the lowest for Chionidae (0.157) and Hydrobatidae (0.166) (Table 4, Figure 2).

**Table 3.** National-scale mean collision risk index for each seabird (sub)family in relation to wind turbines. The risk is expressed as numerical score, with higher values corresponding to a greater potential for collision for the given family.

| **Family** | **Flight**<br>**Altitude** | **Maneuverability** | **Flight Time**<br>**(%)** | **Nocturnal**<br>**Flight** | **Collision**<br>**Risk** |
| --- | --- | --- | --- | --- | --- |
| Chionidae | 0.132 | 0.207 | 0.092 | 0.085 | 0.017 |
| Hydrobatidae | 0.190 | 0.125 | 0.407 | 0.319 | 0.054 |
| Haematopodidae | 0.234 | 0.353 | 0.399 | 0.289 | 0.081 |
| Charadriidae | 0.257 | 0.203 | 0.425 | 0.333 | 0.082 |
| Oceanitidae | 0.224 | 0.187 | 0.602 | 0.414 | 0.090 |
| Phaethontidae | 0.342 | 0.189 | 0.382 | 0.274 | 0.096 |
| Laridae | 0.338 | 0.423 | 0.384 | 0.245 | 0.119 |
| Stercorariidae | 0.422 | 0.270 | 0.475 | 0.249 | 0.140 |
| Scolopacidae | 0.408 | 0.244 | 0.540 | 0.244 | 0.140 |
| Procellariidae | 0.404 | 0.357 | 0.522 | 0.396 | 0.172 |
| Diomedidae | 0.384 | 0.555 | 0.556 | 0.375 | 0.190 |
| Sterninae | 0.501 | 0.256 | 0.568 | 0.317 | 0.190 |
| Sulidae | 0.499 | 0.357 | 0.529 | 0.312 | 0.199 |
| Fregatidae | 0.717 | 0.409 | 0.796 | 0.279 | 0.354 |

**Figure 2.**
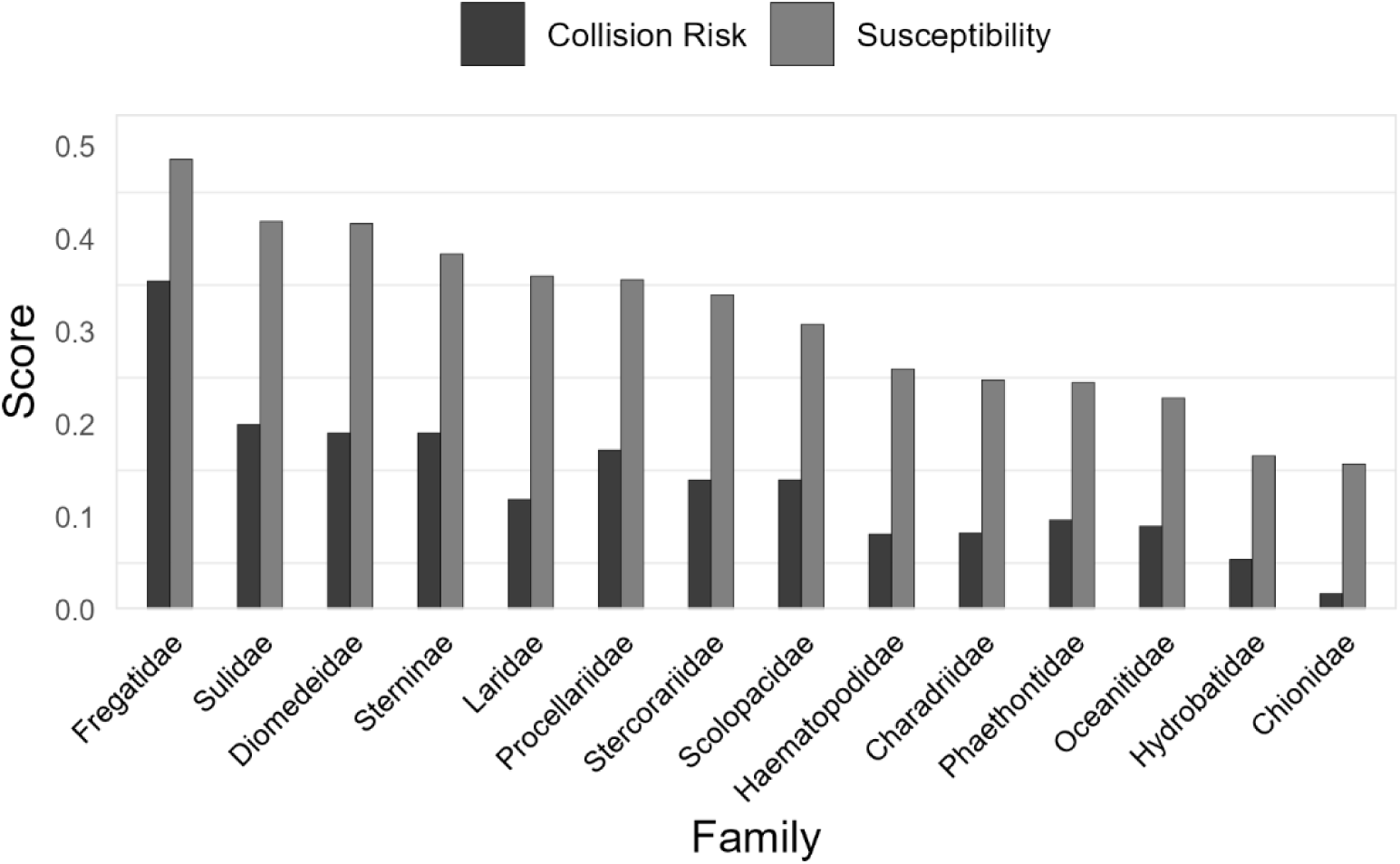
Collision risk and susceptibility indices for each seabird (sub)family. Values are expressed as numerical scores, allowing comparison of relative risk and susceptibility levels across families. Higher scores represent greater potential impact for the given (sub)family.

**Table 4.** National-scale susceptibility index for each seabird (sub)family in relation to wind farms. Susceptibility is expressed as numerical score, with higher values corresponding to a greater susceptibility for the given family.

| **Family** | **Flight**<br>**Altitude** | **Maneuverability** | **Habitat**<br>**specialization** | **Susceptibility** |
| --- | --- | --- | --- | --- |
| Chionidae | 0.132 | 0.207 | 0.264 | 0.157 |
| Hydrobatidae | 0.190 | 0.125 | 0.364 | 0.166 |
| Oceanitidae | 0.224 | 0.187 | 0.547 | 0.228 |
| Phaethontidae | 0.342 | 0.189 | 0.406 | 0.245 |
| Charadriidae | 0.257 | 0.203 | 0.565 | 0.247 |
| Haematopodidae | 0.234 | 0.353 | 0.382 | 0.259 |
| Scolopacidae | 0.408 | 0.244 | 0.541 | 0.307 |
| Stercorariidae | 0.422 | 0.270 | 0.655 | 0.340 |
| Procellariidae | 0.404 | 0.357 | 0.612 | 0.356 |
| Laridae | 0.338 | 0.423 | 0.635 | 0.360 |
| Sterninae | 0.501 | 0.256 | 0.789 | 0.384 |
| Diomedeidae | 0.384 | 0.555 | 0.619 | 0.416 |
| Sulidae | 0.499 | 0.357 | 0.800 | 0.419 |
| Fregatidae | 0.717 | 0.409 | 0.664 | 0.486 |

For the disturbance/displacement species-level index incorporating national conservation status, the species with the highest average scores were *Limnodromus griseus* and *Calidris canutus* (0.098), *Thalasseus maximus* (0.089), *Fregata minor* and *F*. *trinitatis* (0.084) and *Sula sula* (0.082). The lowest scores were observed for *Chionis albus* (0.002) and *Hydrobates leucorhous* (0.008) (Table S15, Figure S1). When using global conservation status, the highest average scores were for *Diomedea dabbenena* (0.092), *Pterodroma incerta* and *Pterodroma madeira* (0.077), and *Thalassarche chlororhynchos* and *Phoebetria fusca* (0.073), while the lowest were again for *C*. *albus* (0.002) followed by *Phaethon aethereus* and *P*. *lepturus* (0.012) (Table S16, Figure S2).

### 3.3 Spatial distribution of seabird vulnerability and offshore wind development

Regarding spatial distribution, the most vulnerable states were identified using citizen science data (Figure S3) for each vulnerability index (Figures 3–6). The susceptibility and collision risk assessments for coastal states highlighted the regions where species exhibit the highest sensitivity to the installation of offshore wind complexes along the Brazilian coast. Accordingly, the most sensitive states were Rio Grande do Sul (23.78; 10.69), followed by Santa Catarina (20.86; 9.31) and Rio de Janeiro (20.44; 9.35) (Table 5, Figures S4–S7).

**Figure 3.**
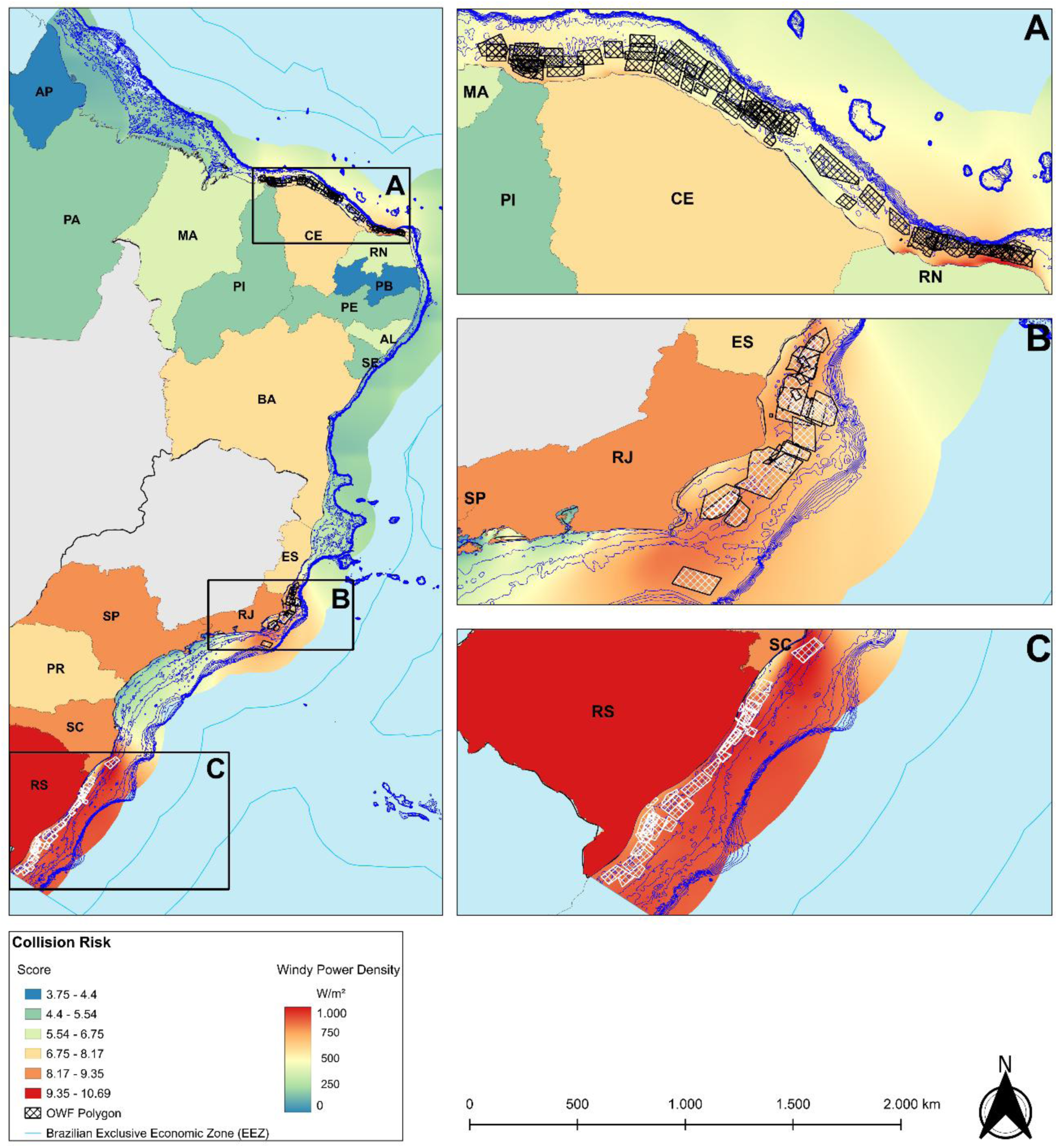
Spatial variation of the seabird collision risk index across Brazilian coastal states. Rectangles indicate focal areas of high vulnerability, highlighting the regional stratification of risk and the need for spatially differentiated environmental licensing for offshore wind farms.

**Figure 4.**
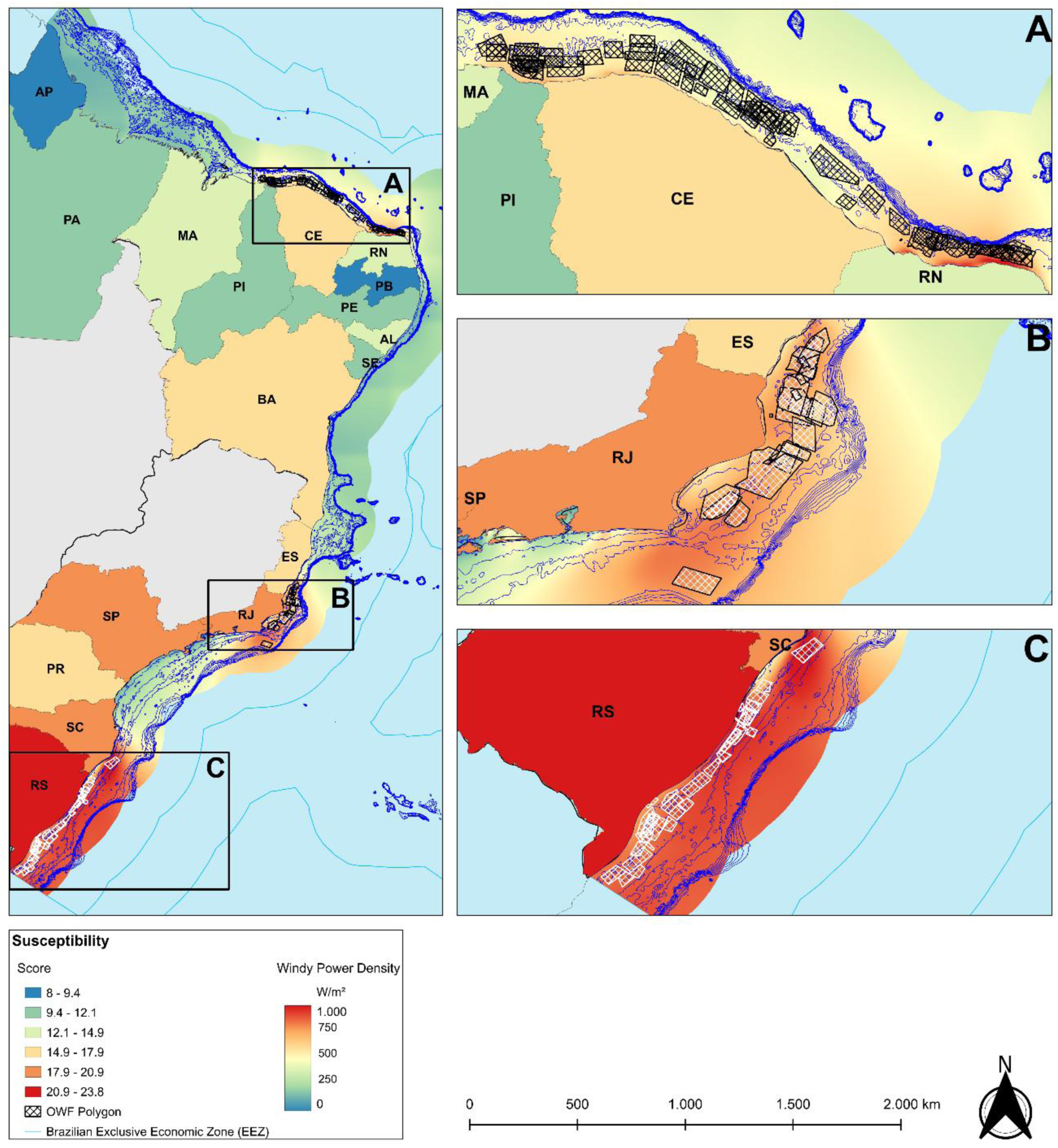
Spatial variation of the seabird susceptibility index across Brazilian coastal states. Rectangles indicate focal areas of high vulnerability, highlighting the regional stratification of susceptibility and the need for spatially differentiated environmental licensing for offshore wind farms.

**Figure 5.**
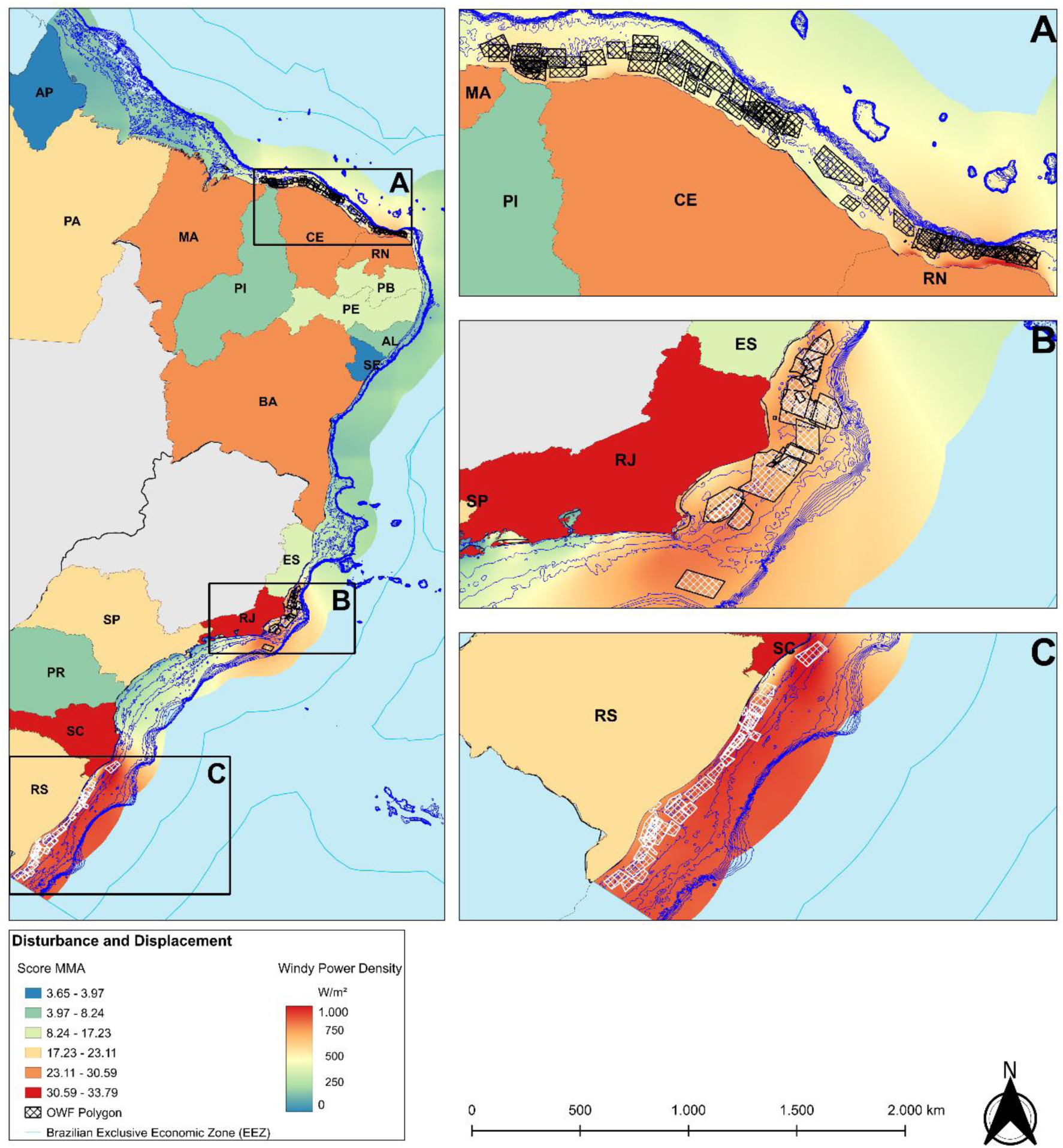
Spatial variation of seabird disturbance and displacement indices according to national conservation status (MMA) across Brazilian coastal states. Rectangles indicate focal areas of high vulnerability, highlighting the regional stratification of disturbance and displacement risks and the need for spatially differentiated environmental licensing for offshore wind farms.

**Figure 6.**
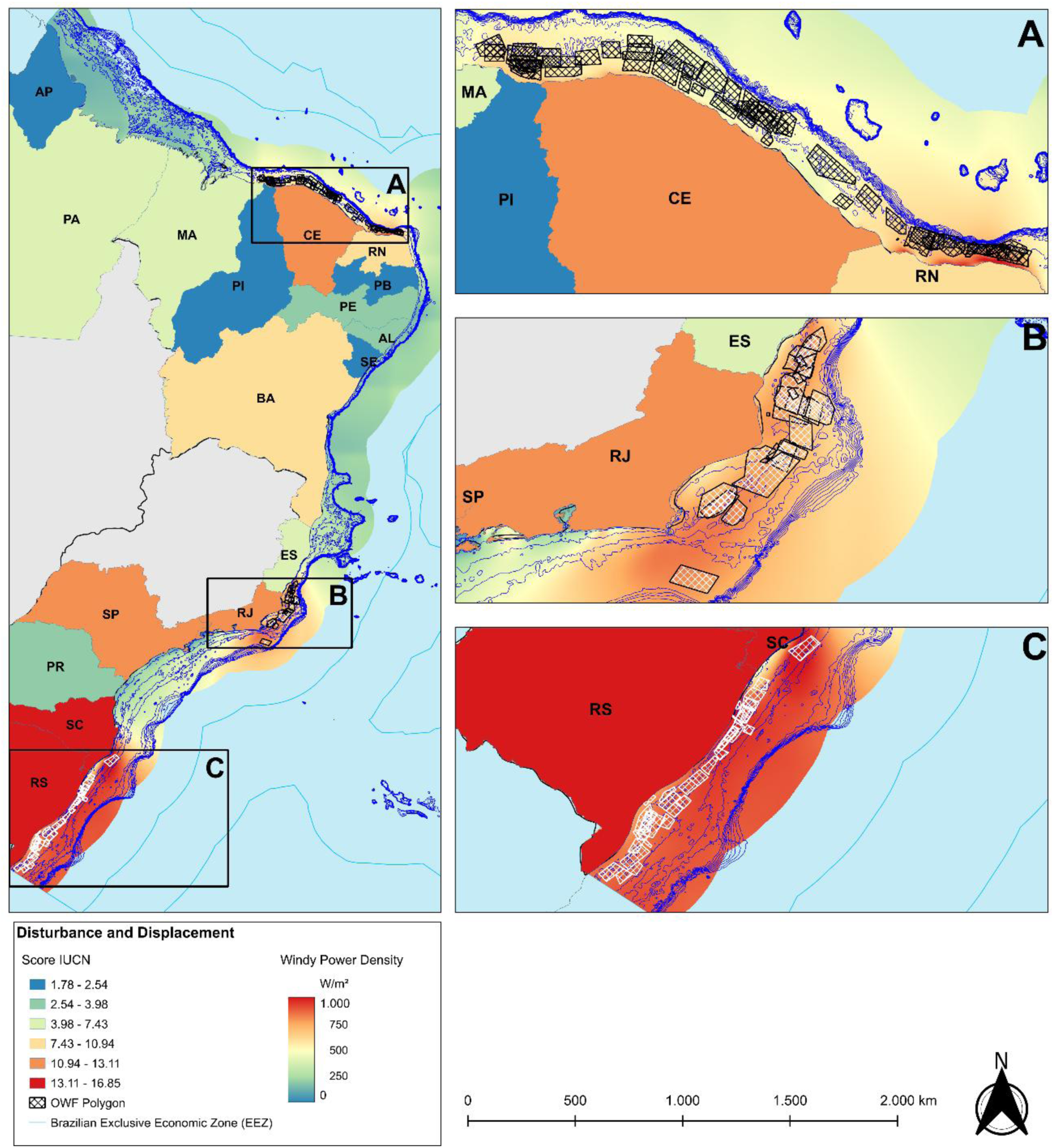
Spatial variation of seabird disturbance and displacement indices according to global conservation status (IUCN Red List) across Brazilian coastal states. Rectangles indicate focal areas of high vulnerability, highlighting the regional stratification of disturbance and displacement risks and the need for spatially differentiated environmental licensing for offshore wind farms

**Table 5.** Susceptibility and collision risk indices for each coastal state. The table presents numerical scores for both indices, where higher values indicate greater susceptibility and greater collision risk for the respective state.

| **States** | **State code** | **Region** | **Susceptibility**<br>**index** | **Collision Risk**<br>**index** |
| --- | --- | --- | --- | --- |
| Paraíba | PB | NE | 8.02 | 3.75 |
| Amapá | AP | N | 9.38 | 4.40 |
| Pernambuco | PE | NE | 10.28 | 4.63 |
| Pará | PA | N | 10.87 | 4.88 |
| Piauí | PI | NE | 11.83 | 5.29 |
| Sergipe | SE | NE | 12.13 | 5.54 |
| Alagoas | AL | NE | 12.93 | 5.90 |
| Maranhão | MA | NE | 14.15 | 6.43 |
| Rio Grande do Norte | RN | NE | 14.94 | 6.75 |
| Ceará | CE | NE | 16.19 | 7.35 |
| Espírito Santo | ES | SE | 17.02 | 7.89 |
| Parana | PR | S | 17.10 | 7.76 |
| Bahia | BA | NE | 17.93 | 8.17 |
| São Paulo | SP | SE | 20.06 | 9.06 |
| Rio de Janeiro | RJ | SE | 20.44 | 9.35 |
| Santa Catarina | SC | S | 20.86 | 9.31 |
| Rio Grande do Sul | RS | S | 23.78 | 10.69 |

## 4. Discussion

This study represents the first national-scale assessment of seabird sensitivity to offshore wind energy development along the Brazilian coast, combining expert elicitation, species trait data, and spatial occurrence records. Our results indicate that certain seabird groups, particularly Fregatidae, Sulidae, and Sterninae, exhibit high susceptibility and collision risk due to their flight behavior and habitat use. At the species level, disturbance and displacement indices varied depending on whether national or global conservation status was applied, underscoring the importance of integrating both perspectives when assessing vulnerability. Spatially, the southern Brazilian coast, especially the state of Rio Grande do Sul, was identified as a high-priority area for impact mitigation, given its high seabird richness and the concentration of planned OWFs projects.

Our findings demonstrate that avian vulnerability to OWFs is not taxonomically homogeneous, as it is directly influenced by a suite of intrinsic biological and behavioral traits specific to each taxonomic group. Species with high maneuverability in flight are expected to suffer less impact from collisions. Conversely, families characterized by predominantly planned flight, low maneuverability, or frequent occupation of the rotor sweep zone are expected to exhibit high vulnerability indices (Fernández-Juricic et al., 2018). Gregarious behavior also acts as a critical factor, with social species being more susceptible to collisions than solitary ones. Accordingly, our study identifies Fregatidae, Sulidae, Sterninae, Procellariidae and Diomedeidae as the groups with the highest vulnerability in Brazil. The extreme collision risk observed for Fregatidae is driven by their reliance on thermal-soaring flight and low aerial maneuverability within the rotor-swept zone. For Sulidae and Sterninae, the risk is exacerbated by foraging strategies such as plunge-diving, which entails a succession of behaviors including patrolling over the sea at moderate flight speeds and altitudes to scan the surface for epipelagic prey (Bugoni and Vooren, 2005). This foraging activity is characterized by a high frequency of flight angle changes, leading to increased exposure time within the rotor-swept zone. This vulnerability is further compounded by frequent daily movements between nesting or roosting sites and foraging grounds. These findings reinforce the global consistency of trait-based assessments, demonstrating that biological characteristics are reliable indicators of vulnerability across distinct marine ecosystems (Cheng et al., 2026).

Studies carried out in Europe have highlighted the high sensitivity of divers (Gaviidae), gannets (Sulidae), and large gulls (Laridae) (Furness et al., 2013; Garthe and Hüppop, 2004), while in the United States (Pacific Coast), brown pelicans (Pelecanidae) and cormorants (Phalacrocoracidae) show the highest collision vulnerability (Adams et al., 2016). In the Southern Hemisphere, Australia exhibits high susceptibility among Procellariidae and Laridae (terns) (Reid et al., 2023). Although Brazil shares certain vulnerable taxa with Australia, such as albatrosses and terns, the prominence of Fregatidae as the most critical group in our study is a hallmark of Brazil’s tropical and subtropical marine environment (Bugoni et al., 2022; Mancini et al., 2016; Reid et al., 2023). Regardless of these regional taxonomic shifts, attributes related to flight height, maneuverability, and foraging ecology remain universal determinants of risk. Identifying these characteristics is fundamental for selecting target species, which should be prioritized as key indicators in environmental monitoring programs from the preliminary licensing phase to the construction and operation phases of offshore wind farms, to ensure ecologically sound, evidence-based, and adaptive management strategies (Croll et al., 2022; Lemos et al., 2023).

The spatial distribution of the most susceptible taxonomic groups is non-uniform along the Brazilian coast, which extends approximately 8,500 km across a latitudinal range from 4.2°N to 33.4°S. This latitudinal breadth facilitates a heterogeneous seascape characterized by the convergence of distinct water masses, localized upwellings, and complex oceanic and shelf fronts (Campos et al., 1995; Piola et al., 2008). The coastline is further structured by the discharge of major river systems, specifically the Amazon and Plata rivers, which create a mosaic of marine and estuarine environments (Daudt et al., 2019; Möller et al., 2008). State-level vulnerability assessments indicate a concentration of risk in the south and southeast regions across all calculated indices, with Rio Grande do Sul, Santa Catarina, and Rio de Janeiro exhibiting the most substantial scores. In the north and northeast regions, vulnerability is associated with the presence of tidal flats and mangroves that serve as critical stopover sites for Nearctic shorebirds, specifically within the families Scolopacidae and Charadriidae (Fedrizzi et al., 2016; Rodrigues, 2000; Siqueira et al., 2024). Furthermore, a recently tracked individual of *F. magnificens* leaving the Fernando de Noronha archipelago was also recorded using the region, which reinforces the importance of environmental planning, considering the extensive home ranges and high spatial connectivity characteristic of this and other species that reproduce in adjacent oceanic archipelagos, in addition to the scarcity of data of this nature (Santos et al., 2025).

The southern region serves as a biological hotspot where high-energy wind zones overlap with the foraging areas of Procellariiformes and Sterninae. While Procellariiformes are distributed across the entire continental shelf, ecological niche modelling indicates that *Thalassarche chlororhynchos*, *Procellaria aequinoctialis* and *Pterodroma incerta* frequently explore inner-shelf coastal waters (Lemos et al., 2023), thereby placing these specific Procellariiformes species alongside Sterninae at the top of the priority list regarding potential spatial conflicts with OWFs within this region. In the Southeast, the high collision risk is primarily driven by the abundance of Fregatidae and Sulidae, as the region hosts the largest Brazilian colonies of both *F. magnificens* and *S. leucogaster* (Alves et al., 2004). The proximity of these breeding sites to potential OWF areas increases the frequency of overlap between the birds’ foraging behaviors, such as thermal-soaring and plunge-diving, and the rotor-swept zone, particularly in the state of Rio de Janeiro. However, the region also harbors important colonies of Sterninae, including species currently listed on the national red list, such as *Thalasseus maximus*, *T. acuflavidus*, and *Sterna hirundinacea*, which are known to perform daily commuting movements between coastal beaches or islands and foraging areas located within the shallow continental shelf (Bugoni et al., 2005; Bugoni and Vooren, 2005; Revorêdo et al., 2025). In this context, the pronounced regional stratification demonstrates that a standardized national framework for offshore wind licensing is insufficient and thus management strategies must be spatially differentiated to address the specific taxonomic sensitivities and oceanographic drivers unique to each coastal sector.

The present study possesses intrinsic limitations arising from necessary methodological choices made in response to Brazil’s geographic scale and data availability. Firstly, taxonomic aggregation at the family level for collision and susceptibility indices, while diverging from Northern Hemisphere approaches which are focused on individual species (Bright et al., 2008; Buckingham et al., 2026; Garthe and Hüppop, 2004), was adopted to circumvent the scarcity of refined biological data for all species within the Brazilian EEZ. Secondly, despite the inclusion of uncertainty estimates to weigh expert inputs, the elicitation method carries inherent subjectivity and potential regional biases. Furthermore, the use of Likert scales may compress continuous biological variations into discrete categories. Additionally, presenting vulnerability at administrative scales (state) rather than through continuous pelagic spatialization reflects the absence of *in situ* density data that would allow for high-resolution static modeling without significant prediction errors for highly mobile species. However, such limitations do not compromise the validity of the results, and they underpin their robustness and strategic necessity. The family-level analysis proved to be an efficient and conservative method, capturing common biological traits and ecological patterns that allow for the recommendation of standardized mitigation measures for groups with shared risk profiles. This scale, when integrated with regional sensitivity, offers a rapid and reliable screening tool for regulatory agencies. Regarding the use of expert knowledge, this approach is scientifically recognized in data-poor scenarios, being the only viable pathway to integrate flight behaviors and seasonal activity variations not yet systematized in national databases (e.g., Tobias et al., 2022).

The reliability of the study is further reinforced by the inclusion of multiple conservation statuses (IUCN and MMA), which prevented the underestimation of risks for critically endangered species at the local level, such as *Limnodromus griseus*. Furthermore, by establishing a national baseline, this work fills a critical gap in the Brazilian Marine Spatial Planning (MSP), recently instituted by Federal Decree No. 12,491/2025. Under this new legal framework, the MSP (locally known as PEM) is mandated to serve as a primary subsidy for environmental licensing processes and a mechanism for ensuring legal certainty by preventing conflicts among maritime stakeholders. By providing a structured, trait-based assessment of avian vulnerability, our study offers the necessary scientific evidence to support the delimitation of management zones and the implementation of regional management plans, ensuring that the expansion of the offshore wind sector within the *Amazônia Azul* aligns with the environmental sustainability goals established by the Brazilian government. Our findings function as a robust prioritization map, indicating where mitigation efforts and investments in high-cost technologies, such as UAV monitoring and biologging of sentinel taxa, should be concentrated. This is especially critical at the project-specific scale, as these indices can assist regulatory agencies, such as IBAMA, in refining environmental diagnosis and monitoring requirements for developers and consultants.

The strategic applicability of this study is direct and essential within the environmental licensing context for OWFs in Brazil, providing the necessary technical support to fulfill IBAMA’s Terms of Reference (TR) requirements. By quantifying collision risk and susceptibility indices for priority seabirds, these results deliver the precise characterization of sensitive groups within the rotor-swept zone required for pre-installation diagnosis. In scenarios where primary tracking or census data are scarce, this assessment establishes an indispensable scientific baseline, allowing developers and consultants to strategically concentrate high-precision efforts on the identified areas and taxa of highest concern. Consequently, the study not only systematizes expert knowledge as per national standards but also provides the framework for marine spatial planning and licensing to transcend a reactive approach, grounding decisions in early evidence of regional sensitivity. Beyond individual licensing, this work plays a fundamental role in strengthening decision-support and strategic energy planning platforms, such as BiodivEPE (developed by Brazil’s Energy Research Office – EPE) and AVISTEP (BirdLife International’s global Avian Sensitivity Tool for Energy Planning). Despite their sophisticated design, both platforms currently lack detailed data layers regarding seabird sensitivity and specific risks in Brazilian waters. By providing quantitative collision and susceptibility indices, this study delivers the scientific content required to bridge this technological gap, allowing these platforms to evolve from species occurrence repositories into genuine risk analysis instruments. Thus, this work establishes itself as a provider of essential data for early screening and safer site selection, ensuring that Brazil’s offshore wind expansion aligns with international best practices for biodiversity conservation. Based on these comprehensive findings and to effectively bridge the gap between scientific evidence and offshore wind management, we propose targeted recommendations below.

## 5. Recommendations

### 5.1 Macro-scale planning and proactive early screening

Regulatory bodies and developers must implement rigorous early screening to completely avoid leasing or building within ecologically sensitive polygons. These include Marine Protected Areas (UCs), Key Biodiversity Areas (KBAs), core foraging hotspots, and critical migratory corridors;

### 5.2 Integration into Marine Spatial Planning

The quantitative vulnerability and risk indices developed in this study should be formally integrated into the layers of the Brazilian Marine Spatial Planning (PEM) framework, recently reinforced by Federal Decree No. 12,491/2025. Utilizing these metrics at the state and national levels allows the PEM to serve as a proactive tool for macro-scale site selection. This transcends reactive, project-by-project licensing and ensures long-term legal certainty for maritime stakeholders within the *Amazônia Azul*;

### 5.3 Cross-scale conservation status integration

Environmental diagnoses must cross-reference both national (MMA) and global (IUCN) conservation statuses. This integration is critical to prevent underestimating the localized extinction risks of specific over-wintering or stopover populations, such as Nearctic shorebirds, which may otherwise be masked by lower global threat categories;

### 5.4 Spatially differentiated licensing frameworks

Regulatory agencies, such as IBAMA, must move away from a standardized, one-size-fits-all national framework for offshore wind licensing. Environmental requirements and mitigation measures must be spatially differentiated to directly address the unique taxonomic sensitivities and oceanographic drivers of each coastal sector. In the South region, emphasis must be placed on high-vulnerability pelagic Procellariiformes utilizing inner-shelf waters, alongside Sterninae displaying high-frequency nearshore movements. In the Southeast region, protocols must prioritize protecting thermal-soaring Fregatidae and plunge-diving Sulidae around major breeding colonies. In the North and Northeast regions, measures must secure coastal tidal flats and mangrove systems critical for resident and migratory shorebirds;

### 5.5 Comprehensive project layout and transmission system integration

Project designs must be adapted during the preliminary licensing phase to minimize spatial overlap with the avian Rotor Swept Zone (RSZ). Developers should optimize turbine spacing, increase lower blade-tip clearance heights, and align layout configurations to preserve local flight pathways. Crucially, the exact positioning of the entire electrical connection system, including subsea cables, land-sea transition corridors across beaches, coastal substations, and overland transmission lines, must be explicitly detailed in the initial project design underpinning the EIA. This comprehensive detailing is imperative to ensure that the cumulative impacts on coastal ecosystems and the specific collision risks for high-wing-load species are evaluated as an integrated system, preventing fragmented licensing. Furthermore, spatial layouts must account for nearby estuarine systems, marine parcéis (shoals), and adjacent oceanic islands, leaving wide, uninhibited corridors to mitigate cumulative, multi-park regional barrier effects;

### 5.6 Three-dimensional farm layout and evasion corridors

When planning the layout, including passageways, the three-dimensional distribution of wind turbines across the site must be designed to minimize obstacles. It is recommended to align the turbines parallel to the primary flight paths of local bird species (rather than perpendicular, which would create a “wall”) and to establish turbine-free corridors that allow birds to move safely between their feeding grounds, thereby avoiding the so-called “barrier effect”;

### 5.7 Integrated Eulerian-Lagrangian field methodologies

Ideally, Environmental Impact Assessments (EIAs) should deploy a combined methodological framework. This includes Eulerian (e.g. systematic ship-based visual censuses and high-definition digital aerial/UAV surveys coupled with automated machine learning) paired with Lagrangian approaches (high-frequency 3D GNSS/GPS biologging). Dataloggers should be equipped with triaxial accelerometers and pressure sensors, particularly for plunge-diving or pursuit-diving sentinel taxa, to segment behaviors and accurately chart fine-scale 3D airspace use;

### 5.8 Robust spatiotemporal BACI and long-term monitoring designs

Studies should strictly adhere to a Before-After Control-Impact (BACI) experimental setup, establishing ecologically comparable reference/control zones outside the Area of Direct Influence (ADA). To account for profound macro-seasonal phenological shifts and high mobile variations, primary baseline diagnostic field surveys must encompass at least one full continuous year. This baseline must be followed by a mandatory, developer-financed post-construction operational monitoring program lasting no less than four years, effectively covering the entire period of the Installation License and the initial years of the Operating License to track long-term micro-, meso-, and macro-scale evasion, attraction (artificial reef effect), or collision dynamics;

### 5.9 Autonomous in-situ automated validation systems

Operational wind farms should be mandated to host autonomous, real-time validation technologies directly on turbine structures. This includes automated fixed and thermal 3D camera tracking systems capable of recording direct collisions or evasion maneuvers during nocturnal flights, foggy periods, or inclement weather. Additionally, automated acoustic sensors (microphones) and fixed automated VHF receiver stations aligned with collaborative international tracking arrays (e.g. Motus network) should be installed to close the regional data gap regarding the coastal connectivity of small migratory species;

### 5.10 Active turbine management, acoustic risk modeling, and data centralization

Active turbine management (shutdown and curtailment), specifically the adoption of technologies to temporarily shut down turbines or reduce blade speed during critical periods, is considered the most efficient way to minimize fatalities during daily or seasonal peak activities and adverse weather conditions. Concurrently, EIAs in the southern continental shelf must include predictive acoustic modeling of underwater noise generated by both pile driving and operational vibrations, focusing on deep-diving taxa like Spheniscidae. To optimize transparency and cumulative impact science, all primary raw biological datasets must be compulsorily submitted to the Biodiversity Data Management System for Environmental Impact Assessment (SISBia) to iteratively feed and recalibrate strategic energy planning and decision-support platforms such as BiodivEPE (EPE) and AVISTEP (BirdLife International).

## Declaration of generative AI and AI-assisted technologies in the writing process

During the preparation of this work the authors used Gemini 3 in order to improve the readability and language of the manuscript. After using this tool/service, the authors reviewed and edited the content as needed and take full responsibility for the content of the published article.

## Data availability

Data will be made available on request.

## Acknowledgements

The authors are deeply grateful to the 23 experts who generously shared their time and specialized knowledge through the elicitation process; their contributions were fundamental to the development of this national baseline. We also thank the thousands of citizen scientists whose records in open-access platforms provided the spatial foundation for our analyses. We thank the Graduate Program in Animal Biology (PPGBAN/UFRGS) and the *Centro de Estudos Costeiros, Limnológicos e Marinhos* (CECLIMAR/UFRGS) for the institutional support and infrastructure provided. This study was supported by the *Conselho Nacional de Desenvolvimento Científico e Tecnológico* (CNPq) through a doctoral scholarship to NH (141829/2023-0). GTN is supported by a research productivity fellowship from the CNPq (312826/2025-7).

